# Sensitivity of bipartite network analyses to incomplete sampling and taxonomic uncertainty

**DOI:** 10.1101/2022.06.28.497912

**Authors:** Cristina Llopis-Belenguer, Juan Antonio Balbuena, Isabel Blasco-Costa, Anssi Karvonen, Volodimir Sarabeev, Jukka Jokela

**Affiliations:** Department of Environmental Systems Science, Institute of Integrative Biology, ETH Zürich, Zürich, Switzerland; Department of Aquatic Ecology, EAWAG, Dübendorf, Switzerland; Cavanilles Institute of Biodiversity and Evolutionary Biology, University of Valencia, Valencia, Spain; Department of Invertebrates, Natural History Museum of Geneva, Geneva, Switzerland; Department of Arctic and Marine Biology, UiT The Arctic University of Norway, Tromsø, Norway; Department of Biological and Environmental Science, University of Jyväskylä, Jyväskylä, Finland; Department of Biology, Zaporizhzhia National University, Zaporizhzhia, Ukraine

**Keywords:** Host-parasite interactions, bipartite networks, sampling issues, sampling completeness, taxonomic resolution, additive effect

## Abstract

Bipartite network analysis is a powerful tool to study the processes structuring interactions in antagonistic ecological communities. In applying the method, we assume that the sampled interactions provide an accurate representation of the actual community. However, acquiring a representative sample may be difficult as not all species are equally abundant or easily identifiable. Two potential sampling issues can compromise the conclusions of bipartite network analyses: failure to capture the full range of interactions of species (sampling completeness) and failure to identify species correctly (taxonomic resolution). These sampling issues are likely to co-occur in community ecology studies. We asked how commonly used descriptors (modularity, nestedness, connectance and specialisation (H_2_′)) of bipartite communities are affected by reduced host sampling completeness, parasite taxonomic resolution and their crossed effect. We used a quantitative niche model to generate replicates of simulated weighted bipartite networks that resembled natural host-parasite communities. The combination of both sampling issues had an additive effect on modularity and nestedness. The descriptors were more sensitive to uncertainty in parasite taxonomic resolution than to host sampling completeness. All descriptors in communities capturing less than 70% of correct taxonomic resolution strongly differed from correctly identified communities. When only 10% of parasite taxonomic resolution was retained, modularity and specialisation decreased ∼0.3 and ∼0.1-fold respectively, and nestedness and connectance changed ∼0.7 and ∼3.2-fold respectively. The loss of taxonomic resolution made the confidence intervals of estimates wider. Reduced taxonomic resolution led to smaller size of the communities, which emphasised the larger relative effect of taxonomic resolution on smaller communities. With regards to host sampling completeness, connectance and specialisation were robust, nestedness was reasonably robust (∼0.2-fold overestimation), and modularity was sensitive (∼0.5-fold underestimation). Nonetheless, most of the communities with low resolution for both sampling issues were structurally equivalent to correctly sampled communities (i.e., more modular and less nested than random assemblages). Therefore, modularity and nestedness were useful as categorical rather than quantitative descriptors of communities affected by sampling issues. We recommend evaluating both sampling completeness and taxonomic certainty when conducting bipartite network analyses. We also advise to apply the most robust descriptors in circumstances of unavoidable sampling issues.

**Open Research statement:** we provide permanent and open access links to data sources and replication code in Appendix S1.

## Introduction

Ecological and evolutionary processes occur in networks of interacting species. Species interactions are diverse, numerous and often asymmetric due to the unequal dependence between the interacting species (Dormann et al. 2017). These attributes make ecological networks of interactions complex, hampering our ability to disentangle ecological and evolutionary dynamics and to understand responses to changing environments. Despite the complexity of ecological communities, bipartite network analysis has allowed researchers to tackle fundamental research questions and give advice on biodiversity management (Delmas et al. 2019).

Bipartite network analysis is based on the assessment of the distribution of interactions between nodes of different guilds (Blüthgen 2010). For example, in host-parasite bipartite networks, host and parasite species are nodes. Bipartite network analysis does not consider interactions among nodes of the same type. For example, parasite-parasite or host-host interactions are not analysed. Therefore, to make ecological communities tractable, bipartite network analysis assumes that inter-guild interactions are more relevant for ecological communities than intra-guild interactions (Poulin 2010). Bipartite interactions can be weighted (as opposed to simple presence-absence of interactions) to capture a quantitative description of the processes in the natural environment. For example, host-parasite interactions can be weighted by the abundance of each parasite species in each host (Cardoso et al. 2021).

The study of ecological networks not only relies on recording the species composition but also on obtaining large enough samples to build a fair representation of the interactions in communities (Henriksen et al. 2019). Representative samples of ecological communities can be difficult to obtain due to community diversity (Chacoff et al. 2012), temporal and spatial variation in community structure (Poisot et al. 2015), different level of population and community fragmentation (Frankham et al. 2017) or differences in interaction patterns among individuals of a given species (Guimarães Jr. 2020). Insufficient sampling of both species and their interactions makes it difficult to tease apart biological processes from methodological artifacts. Sampling issues may therefore influence the observed network properties and the conclusions extracted from them (Vizentin-Bugoni et al. 2016).

It is assumed that a sample is a good representative of the actual community when species richness, interaction richness or even network descriptors reach an asymptote (Henriksen et al. 2019). However, acquiring an asymptotic sample of the interactions in a community requires a higher sampling effort than estimating species richness because there are more combinations of pairwise interactions than species (Henriksen et al. 2019). In addition, common interactions of abundant species are detected with low effort, but high sampling effort is required to record rare interactions of less abundant species (Chacoff et al. 2012, Henriksen et al. 2019).

Another confounding factor for bipartite network analysis is the reliability of species identification (Thompson and Townsend 2000). Although, taxonomic accuracy and community size usually represent a trade-off (Renaud et al. 2020), small networks may be more affected by inappropriate identification of a species since it would represent a higher proportion of taxa than in a large network. Taxonomic resolution can be variable both within and between communities, which complicates comparative studies. Often, some nodes are identified as species, whereas other nodes aggregate coarser taxonomic categories or entities within a community (e.g., detritus) or include hidden diversity (e.g., cryptic species) (Thompson and Townsend 2000). Furthermore, loss of taxonomic resolution was found to have a higher impact on the predicted structure of antagonistic insect-plant networks than on their mutualistic counterparts due to the closer dependence of the consumers on their resources in the former type (Rodrigues and Boscolo 2020). Opposite to low taxonomic resolution is species over-splitting, that is identifying individuals of the same species as different species due to phenotypic or other differences (Isaac et al. 2004).

Finally, these sampling issues occur independently and may coincide in the same dataset. For instance, an ecological dataset may present poor sampling completeness regardless of the taxonomic resolution of the few sampled taxa. Then, researchers must strive to control all potential sampling issues at the same time to ensure a correct representation of an ecosystem. Even though sampling issues are known to affect bipartite network descriptors (Rivera-Hutinel et al. 2012, Rodrigues and Boscolo 2020) and may simultaneously affect the same survey, we do not yet know how the combined effect of sampling issues can mislead the interpretation of the structure of ecological communities. We will address this issue by focussing on host-parasite bipartite networks. However, since sampling issues can occur in any community study, our approach is not restricted to host-parasite interactions and can be applied to any bipartite symbiotic interactions.

Parasitic life-history strategies are spread across the whole tree of life. Parasites are present in all ecosystems and dominate diversity in terms of richness and abundance of species, and biomass (Carlson et al. 2020). This astonishing diversity makes host-parasite associations one of the most common types of interactions in ecological communities (Lafferty et al. 2006). Bipartite network analysis has been a key method for research in evolutionary ecology of host-parasite interactions over the last decades (Runghen et al. 2021), including epidemiological and public health issues (Bellekom et al. 2021).

Host individuals are typically the sampling units in ecological and evolutionary parasitology. Commonly, data from host individuals of the same species are pooled together to obtain the parasite community of each host species (Morand and Krasnov 2008). As more host individuals are sampled in a community, the probability of finding an unrecorded parasite species or a new host-parasite interaction reduces, and parasite species richness or number of species interactions approaches an asymptote (Henriksen et al. 2019). At the same time, studies listing parasite species are often characterised by poor taxonomic resolution at least for certain taxa. Such inaccurate assessments of parasite diversity seriously hamper our ability to understand host-parasite dynamics (Poulin and Leung 2010) since it decreases the variability in the interaction pattern of different species (Delmas et al. 2019).

Our goal was to understand how decreasing gradients of host sampling completeness, parasite taxonomic resolution and their crossed effect affect four commonly-used descriptors of host-parasite communities: modularity, nestedness, connectance and specialisation (H_2_′) (Table 1). We generated replicates of a simulated host-parasite community, which are not usual in empirical datasets. We resampled the replicate simulated communities controlling for both sampling issues. First, we gradually removed host-parasite interactions to infer the effect of decreasing host sampling completeness on the four descriptors. Second, we evaluated how the four descriptors were influenced by decreasing resolution of parasite taxonomy. To do so, we gradually reduced the number of parasite species in the communities by hierarchically lumping their interactions according to their overlap in host use, which resembles closely related cryptic species that have ecologically similar requirements. Finally, we evaluated the crossed effect of both sampling issues as they are likely to occur simultaneously.

**Table 1.**
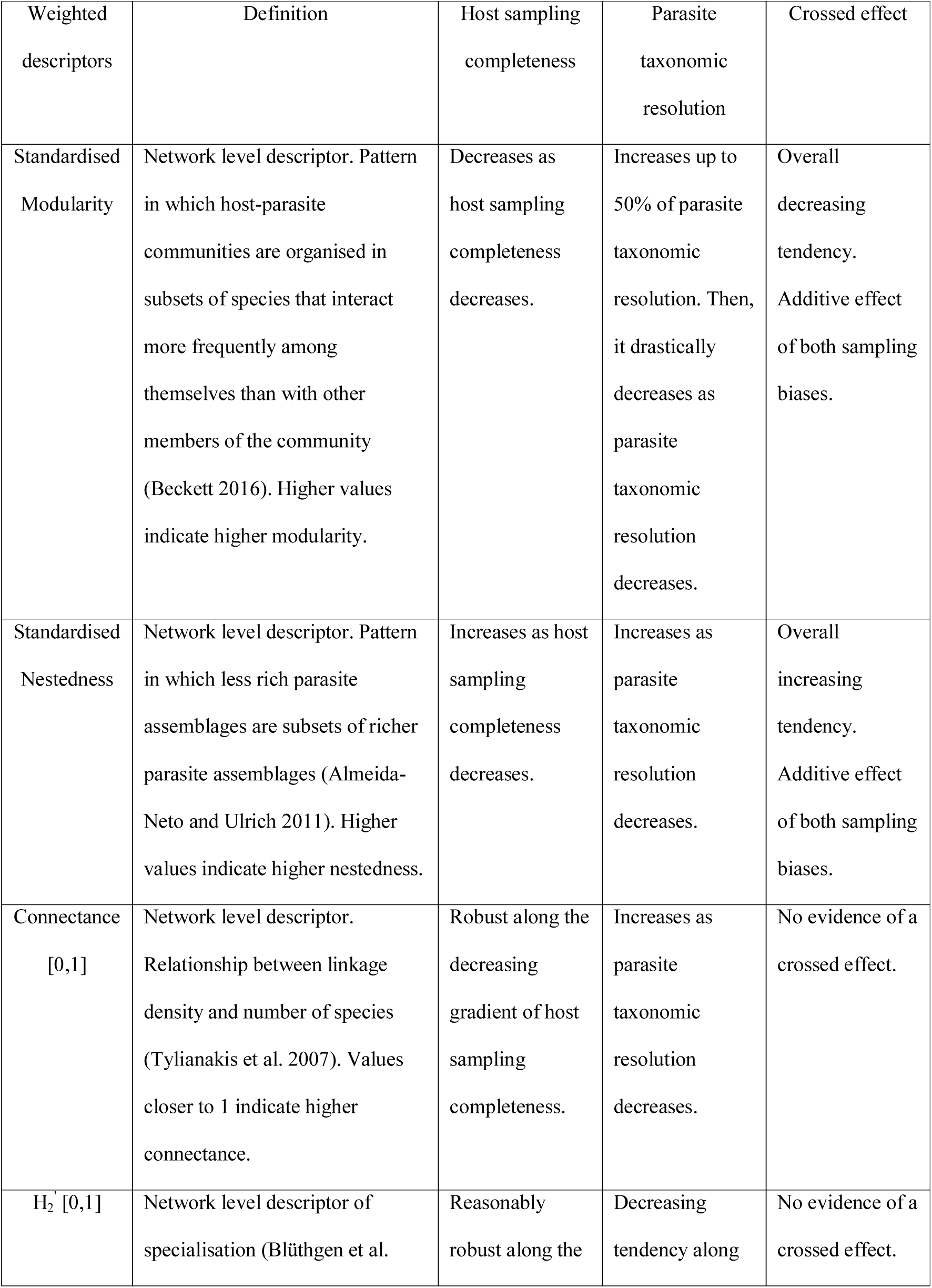
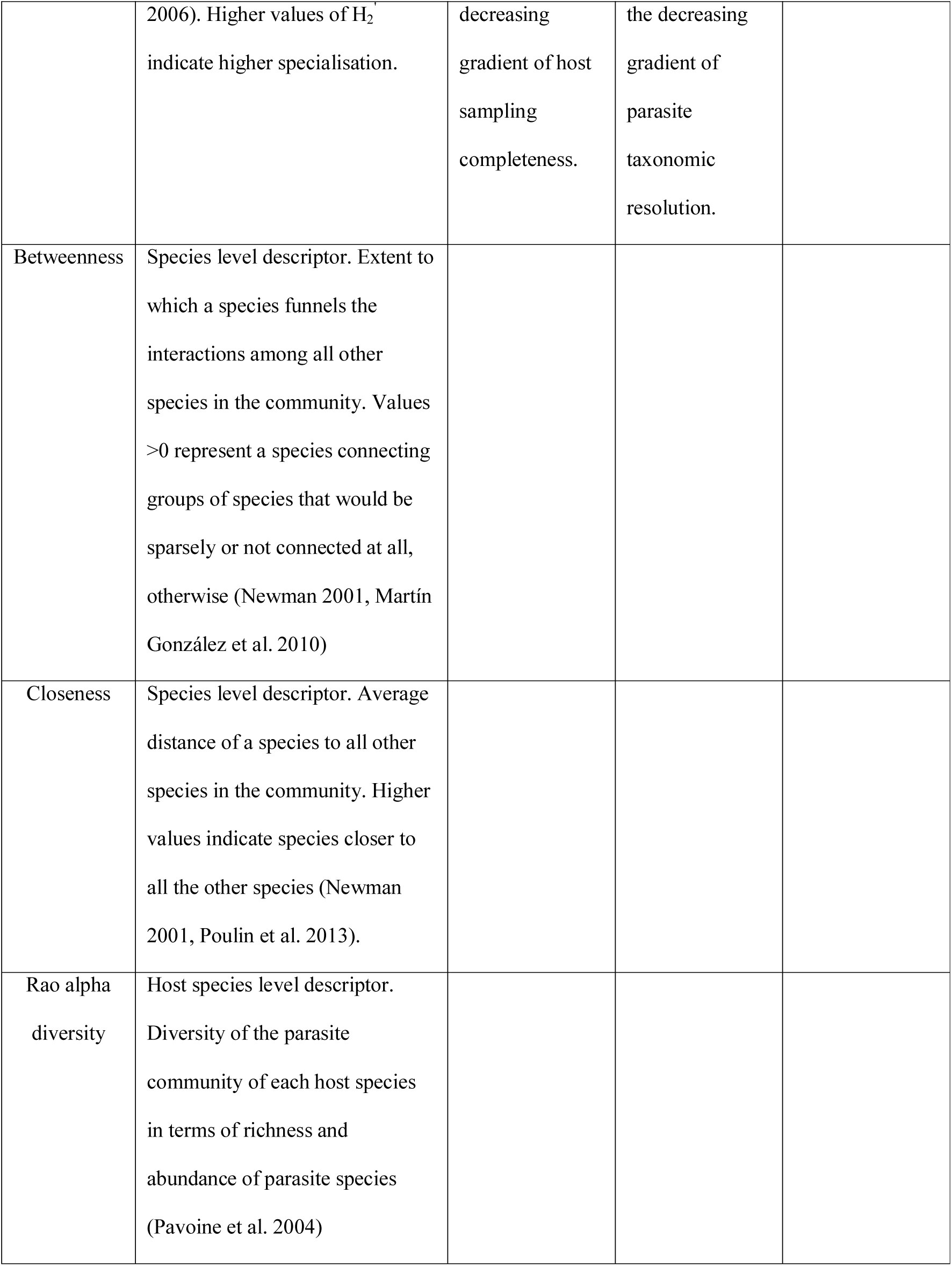
Effect of sampling issues on community and species level descriptors.

We hypothesised that the descriptors would be more robust to host sampling completeness than to parasite taxonomic resolution, at least in communities that are not severely affected by sampling issues (Hypothesis 1). Host individuals of the same species commonly sustain similar parasite communities and, in this sense, represent biological replicates of the same system (Llopis-Belenguer et al. 2020). Therefore, when decreasing host sampling completeness, the overall infection pattern of the community is maintained until the sample size reaches a threshold below which no inference is supported. On the contrary, we expected the reduction in parasite taxonomic resolution to have a greater impact on the community descriptors. Commonly, parasite species are not redundant in their host ranges. Specialisation in their host limits the ability of parasite species to infect many host species (i.e., environmental filtering) (Llopis-Belenguer et al. 2020). Therefore, the detection of the infection patterns would be more sensitive to taxonomic resolution following the loss of inter-species variability. In addition, we expected the combined effect of both sampling issues to cause a greater bias in the descriptors than either of the simulated sampling biases in isolation (Hypothesis 2). Finally, we also hypothesised that the four network descriptors would differ in sensitivity (Hypothesis 3). Based on the results of previous studies on ecological systems outside the host-parasite realm (Blüthgen et al. 2006, Vizentin-Bugoni et al. 2016), redundant interactions would make the descriptors considerably robust against reduced sampling completeness. With regard to taxonomic resolution (Thompson and Townsend 2000, Rodrigues and Boscolo 2020), modularity, nestedness and specialisation would be reasonably robust to loss of taxonomic resolution. However, communities in the gradient of parasite taxonomic resolution varied in size. Connectance expresses the proportion of realised interactions out of all possible interactions (Table 1). A realised interaction represents a higher proportion in a small network than in a larger network. For the same number of interactions, communities with low parasite taxonomic resolution (small networks) would present higher connectance than communities with correct taxonomic resolution (large networks). Then, we expected connectance to be sensitive to parasite taxonomic resolution.

## Material and Methods

### Building simulated communities

Our main dataset consists of 10 replicate simulated networks (hereafter, “full communities”) that were constructed using host-parasite community parameters extracted from published host-parasite community data (n = 6, “natural communities”) (Fig. 1a, Appendix S1). To build the full communities, we used a quantitative niche model (see below) that was initiated with the mean number of host and parasite species (nhost = 13; npara = 42) of the six natural communities. The mean number of interactions per parasite species in the full communities (maxobs.rf = 2,067) was the mean overall number of interactions in the six natural communities (ni = 86,794) divided by npara. In other words, these 10 replicate full communities represent a generalisation of different natural host-parasite communities with respect to their number of species and their number of interactions (Fig. 1a). We performed all the analyses in R (R Core Team 2021). If not specified otherwise, the functions mentioned below are available in bipartite package (Dormann 2011). Replication code and data are provided in Llopis-Belenguer et al. (to be published in Zenodo after acceptance).

**Figure 1.**
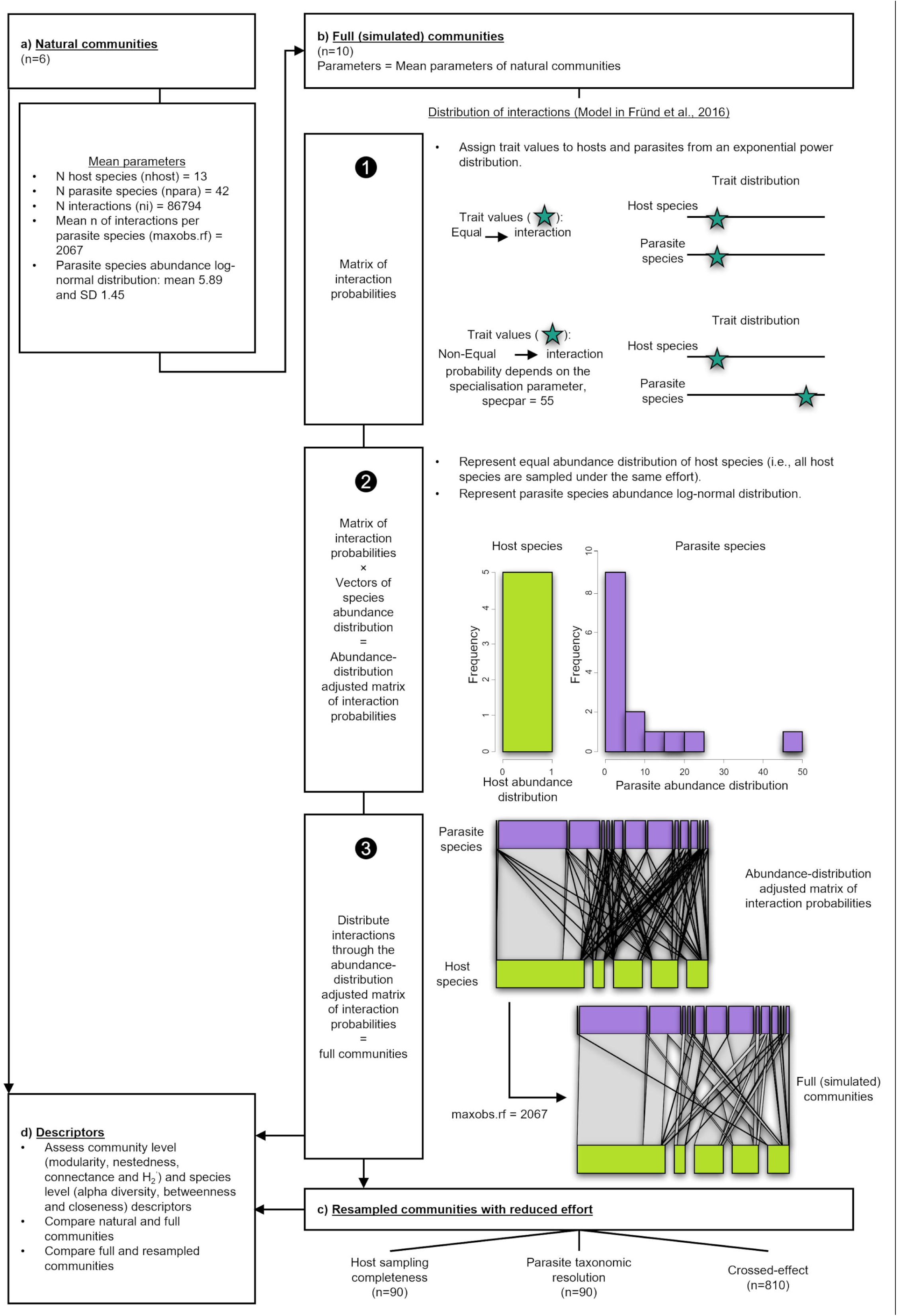
Analysis steps. a) Obtain mean parameters of the natural communities to fit the model. b) Create host-parasite simulated communities (full communities). (1) Build a matrix of interaction probabilities by trait matching and according to a specialisation parameter. (2) Adjust the matrix of interactions probabilities by species abundance distributions. (3) Distribute interactions through the abundance adjusted matrix of interaction probabilities (Fründ et al. 2016). c) Simulate nine levels of sampling biases for host sampling completeness and parasite taxonomic resolution, and their crossed effect. d) Assess and compare network descriptors between (1) natural and full communities and (2) full and resampled communities.

We used the quantitative niche model described in Fründ et al. (2016) to create the full communities (Fig. 1b). The model generates weighted bipartite networks reflecting a chosen specialisation parameter (Fründ et al. 2016). First, the model creates a matrix of interaction probabilities based on quantitative trait values of each host and parasite species (Fig. 1b: 1. Matrix of interaction probabilities). Equal trait values of a host and a parasite represent a realised interaction. Non-equal trait values receive an interaction probability depending on the specialisation parameter (Fründ et al. 2016), which defines the shape (width) of the gaussian niche function. In our case, we assigned trait values to host and parasite species from an exponential power distribution, since the expected distribution of key traits (e.g., body mass) for interaction establishment is usually skewed (Poulin and Morand 1997, Kozłowski and Gawelczyk 2002). We used the highest specialisation parameter used in Fründ et al. (2016) (specpar = 55) (Appendix S2) because specialisation is the general trend in metazoan host-parasite interactions (Poulin 2007). We implemented this procedure with the function *makeweb* (Fründ et al. 2016). Second, we multiplied the interaction probability matrices by species abundance-distribution vectors with the function *make_trueweb* (Fründ et al. 2016) (Fig. 1b: 2. Abundance-distribution adjusted matrix of interaction probabilities). This step adjusts the interaction probability matrix according to the relative abundance of each species (Fründ et al. 2016), i.e. it considers that it is more likely to record interactions between two abundant species than between two rare ones. We assumed an even abundance distribution of the host species. This corresponds to sampling procedures where an effort is made to capture the same number of individuals of each host species (Poulin 1998). We distributed the abundance of parasite species from a log-normal distribution with mean 5.89 and SD 1.45 both in the log-scale, which are similar to the mean and SD of parasite species in the natural host-parasite communities, with the function *get_skewedabuns* (Fründ et al. 2016). Therefore, only the abundance distribution of parasite species affected the interaction probability in our simulation. As the last step, we weighted the full communities with parasite abundances according to the given interaction probability matrix with the function *sampleweb* (Fründ et al. 2016) (Fig. 1b: 3. Full communities). We assumed a mean number of interactions per parasite species equal to maxobs.rf.

### Simulating sampling completeness and taxonomic resolution biases

We resampled the full communities with a reduced effort to simulate communities along decreasing gradients of host sampling completeness, parasite taxonomic resolution and their crossed effect (hereafter, “resampled communities”) (Fig. 1c). Each full community was resampled in 10% steps from 90% to 10% of host sampling completeness and parasite taxonomic resolution, thus simulating a situation where researchers do not have a-priori knowledge of the true size and taxonomic structure of the communities they are sampling.

To simulate the decreasing gradient of host sampling completeness, we used data on parasite abundance in 3,258 fish host individuals from 51 locations and of 41 species (63.9 ±71.4 fish individuals per species) (Appendix S1). Only one of these datasets belongs to the six natural communities we used to derive the full communities (Valtonen et al. 2001). Fish individuals of each fish species and location formed an independent dataset. We resampled host individuals of each independent dataset by reducing the sample size in 10% steps starting from 90% and ending with 10% of the original host individuals. Then, we quantified the mean percentage of interactions per parasite species remaining in all datasets (Table 2a). We applied these resampled percentages to the mean number of interactions per parasite species (maxobs.rf = 2,067) to fill the existing matrix of interaction probabilities in each of the 10-replicate full communities. Thus, we simulated the effect of a decreasing number of sampled host individuals by reducing the mean number of interactions per parasite species in the full communities (Table 2a). We derived 90 resampled communities to examine the effect of host sampling completeness.

**Table 2.** Decreasing gradients of (a) host sampling completeness and (b) parasite taxonomic resolution.

|  | Full communities | Resampled communities |  |  |  |  |  |  |  |  |
| --- | --- | --- | --- | --- | --- | --- | --- | --- | --- | --- |
| (a) Host sampling completeness |  |  |  |  |  |  |  |  |  |  |
| % of remaining host individuals | 100 | 90 | 80 | 70 | 60 | 50 | 40 | 30 | 20 | 10 |
| % of remaining interactions | 100 | 85.1 | 78.5 | 69.3 | 57.2 | 50.5 | 36.8 | 27.7 | 17.9 | 8.2 |
| Mean number interactions per parasite species | 2067 | 1759 | 1622 | 1433 | 1182 | 1043 | 761 | 574 | 371 | 170 |
| (b) Parasite taxonomic resolution |  |  |  |  |  |  |  |  |  |  |
| % of parasite species | 100 | 90 | 80 | 70 | 60 | 50 | 40 | 30 | 20 | 10 |
| Remaining number of parasite species | 42 | 37 | 33 | 29 | 25 | 21 | 16 | 12 | 8 | 4 |

To simulate the effect of incomplete parasite taxonomic resolution, we calculated pairwise Jaccard similarities between all parasite species based on their host-range overlap (Martinez 1991). These pairwise distance estimates were used to distribute the species to a number of groups according to the decreasing gradient of parasite taxonomic resolution (Table 2b). Similar parasite species in the Jaccard calculation were assigned to the same group. We implemented the “around medoids” clustering, which is a more robust version of “k-means” clustering, to distribute parasites into groups. This was performed with the function *clara* in cluster package (Maechler et al. 2021). For example, in the simulation of “50%” of parasite taxonomic resolution, we asked the function to assign parasite species to 21 groups, which represents 50% of the original number of parasite species in the full communities (npara). Consequently, we clustered parasite species based on the similarity of their host range, which resembles the taxonomic resolution issues common in the parasitological data given that closely related species may share hosts.

Table 2b shows the number of groups for each decreasing resolution class. Note that the overall number of interactions (ni) was kept unchanged by this regrouping of parasite species to fewer groups. For each replicate full community, we ran a new clustering algorithm using the same target number of groups (Table 2b). We obtained 90 resampled communities to examine the effect of parasite taxonomic resolution.

Finally, we simulated the crossed effect of host sampling completeness and parasite taxonomic resolution. A gradient of parasite taxonomic resolution (as described above) was created for each resampled community used to examine host sampling completeness. This produced 810 additional resampled communities, which were biased for both sampling issues to varying degrees.

### Community and species level descriptors

We assessed four weighted community level descriptors for each of the 1,000 full and resampled communities: modularity, nestedness, connectance and specialisation (H_2_′) (Table 1). Specifically, we used the Becket algorithm (Beckett 2016) to calculate modularity in our communities with the function *computeModules*. The function *networklevel* was used to measure the weighted versions of the algorithms: NODF (Nestedness metric based on Overlap and Decreasing Fill, Almeida-Neto & Ulrich, 2011), connectance (Tylianakis et al. 2007) and H_2_′ (Blüthgen et al. 2006). We standardised modularity and nestedness of each web since raw values of these descriptors are not directly comparable. To do this, we created 1,000 null communities for each of the 1,000 full and resampled communities with the function *vaznull* (Vázquez et al. 2007). This algorithm creates null host-parasite communities by randomising the total number of host-parasite interactions observed in the full communities. Thus it constrains connectance but not the marginal totals (Vázquez et al. 2007). We then calculated the mean value and the standard deviation of modularity and nestedness of each set of 1,000 null communities. Finally, we standardised modularity and nestedness of each full and resampled community following the equation of the Standardised Effect Size (SES) (Gotelli and Rohde 2002) (Equation 1):

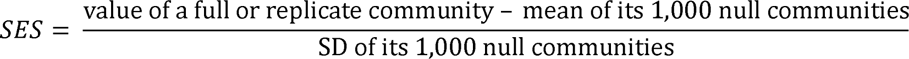

Finally, to assess the effect of each sampling issue and their crossed effect on the community descriptors, four two-way ANOVAs were used with host sampling completeness and parasite taxonomic resolution as fixed factors. If the two fixed factors are significant, a significant interaction term indicates a synergistic effect of both sampling issues, otherwise additive (Ferguson and Stiling 1996).

We used t-tests to establish whether network descriptors of the simulated full communities differed significantly from those of natural communities. Additionally, we aimed to know whether the full communities reproduced the pattern of interactions of the natural communities at the species level. For each full and natural community, we computed Rao alpha diversity of the parasite communities of each host species, and two species-level network descriptors of centrality for both host and parasite species: weighted betweenness and weighted closeness (Table 1). Rao alpha diversity, which accounts for the richness and abundance of species (Pavoine et al. 2004), was calculated with the function *dpcoa* in ade4 package (Thioulouse et al. 2018). To make alpha diversity comparable across host species, we transformed alpha diversity results into their equivalent numbers (E_alpha_) following Equation 2 (Ricotta and Szeidl 2009):

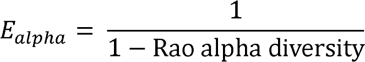

We used a linear regression of E_alpha_ on the communities followed by ANOVA to test differences in E_alpha_ between the full and natural communities. Finally, the function *specieslevel* was used to compute the two descriptors of centrality, betweenness and closeness (Table 1). Both are based on the Dijkstra’s algorithm to find the shortest path in networks of interactions (Newman 2001). We ran the analyses of betweenness and closeness for host and parasite species independently. We excluded those species that were not linked to any other species in the community since no centrality measure could be calculated. To compare these descriptors across communities independently of their sizes, we first regressed each descriptor on their community size. We used the residuals of these regressions as the dependent variables in linear models as a function of the type of study (natural vs simulated) followed by ANOVA to test the differences between full and natural communities (Morris et al. 2014).

## Results

Community and species level descriptors of the full communities did not significantly differ from descriptors of the natural communities, except for closeness (Table 3).

**Table 3.** Comparisons of the weighted descriptors between the full and natural communities. M: standardised modularity. N: standardised nestedness. C: connectance. H_2_’: specialisation. B: betweenness. Cl: closeness. p: parasites. h: hosts.

|  |  | Community level |  |  |  | Species level |  |  |  |  |
| --- | --- | --- | --- | --- | --- | --- | --- | --- | --- | --- |
|  |  | M | N | C | H<sub>2</sub>' | Bp | Bh | Clp | Clh | Rao<br>alpha<br>diversity |
| Full<br>communities<br>(n=10) | Minimum | 54.32 | -14.03 | 0.05 | 0.67 | 0 | 0 | 0.001 | 0.02 | 1.08 |
|  | Maximum | 77.80 | -10.59 | 0.07 | 0.84 | 0.46 | 0.46 | 0.13 | 0.22 | 7.9 |
|  | Mean | 66.25 | -11.90 | 0.05 | 0.76 | 0.02 | 0.08 | 0.02 | 0.09 | 3.22 |
|  | Standardised<br>error | 2.30 | 0.30 | 0.00 | 0.01 | 0.003 | 0.01 | 0.001 | 0.00<br>4 | 0.12 |
| Natural<br>communities<br>(n=6) | Minimum | 36.93 | -13.39 | 0.04 | 0.44 | 0 | 0 | 10<sup>-5</sup> | 0 | 1 |
|  | Maximum | 150.2<br>1 | -8.67 | 0.07 | 0.88 | 0.77 | 0.72 | 0.08 | 0.7 | 9.54 |
|  | Mean | 74.76 | -11.09 | 0.05 | 0.66 | 0.02 | 0.08 | 0.005 | 0.06 | 2.73 |
|  | Standardised<br>error | 17.47 | 0.71 | 0.00 | 0.06 | 0.005 | 0.02 | 6×10<sup>-4</sup> | 0.01 | 0.19 |
| t-test<br>(community<br>level)<br>ANOVA<br>(species<br>level) | p-value | 0.65 | 0.33 | 0.92 | 0.16 | 0.33 | 0.88 | 0.00 | 0.00 | 0.24 |

Standardised modularity and nestedness showed opposite patterns with respect to host sampling completeness (Fig. 2a and Fig. 2b, darkest purple points and lines). While modularity decreased with decreasing host sampling completeness (∼0.5-fold), nestedness increased (∼0.2-fold). Estimates of modularity in simulations capturing 70% or less of host sampling completeness strongly differed from those values of the full communities (Fig. 2a, error bars do not overlap). Estimates of nestedness in simulations from 40% of host sampling completeness and below strongly differed from estimates of the full communities (Fig. 2b, error bars do not overlap). In contrast, weighted connectance and H_2_′ were robust to host sampling completeness since resampled communities showed little evidence of difference from estimates for full communities (Fig. 1c-d, Table 4).

**Figure 2.**
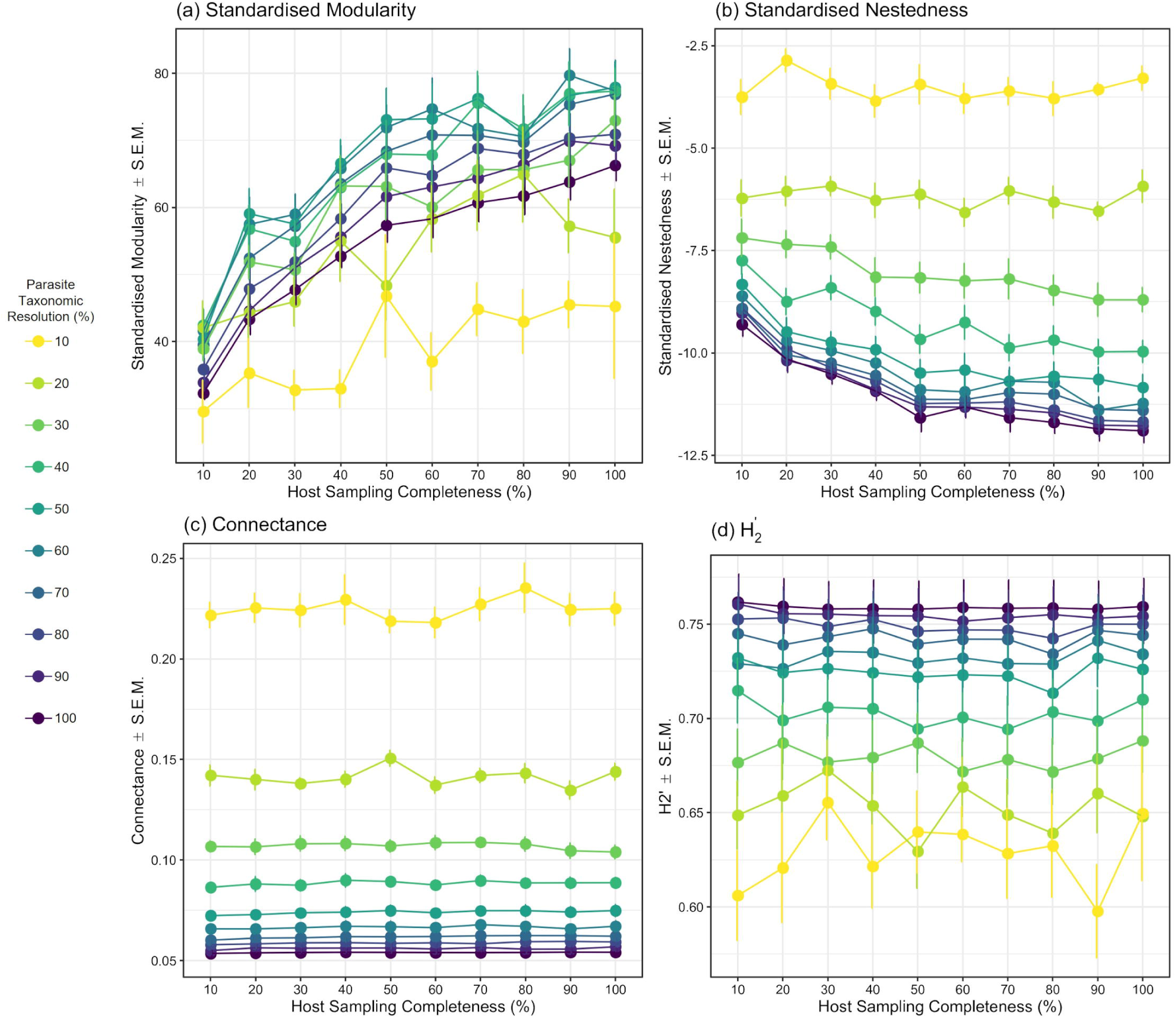
Effect of the decreasing gradient of host sampling completeness, parasite taxonomic resolution and their crossed effect on host-parasite community descriptors.

**Table 4.**
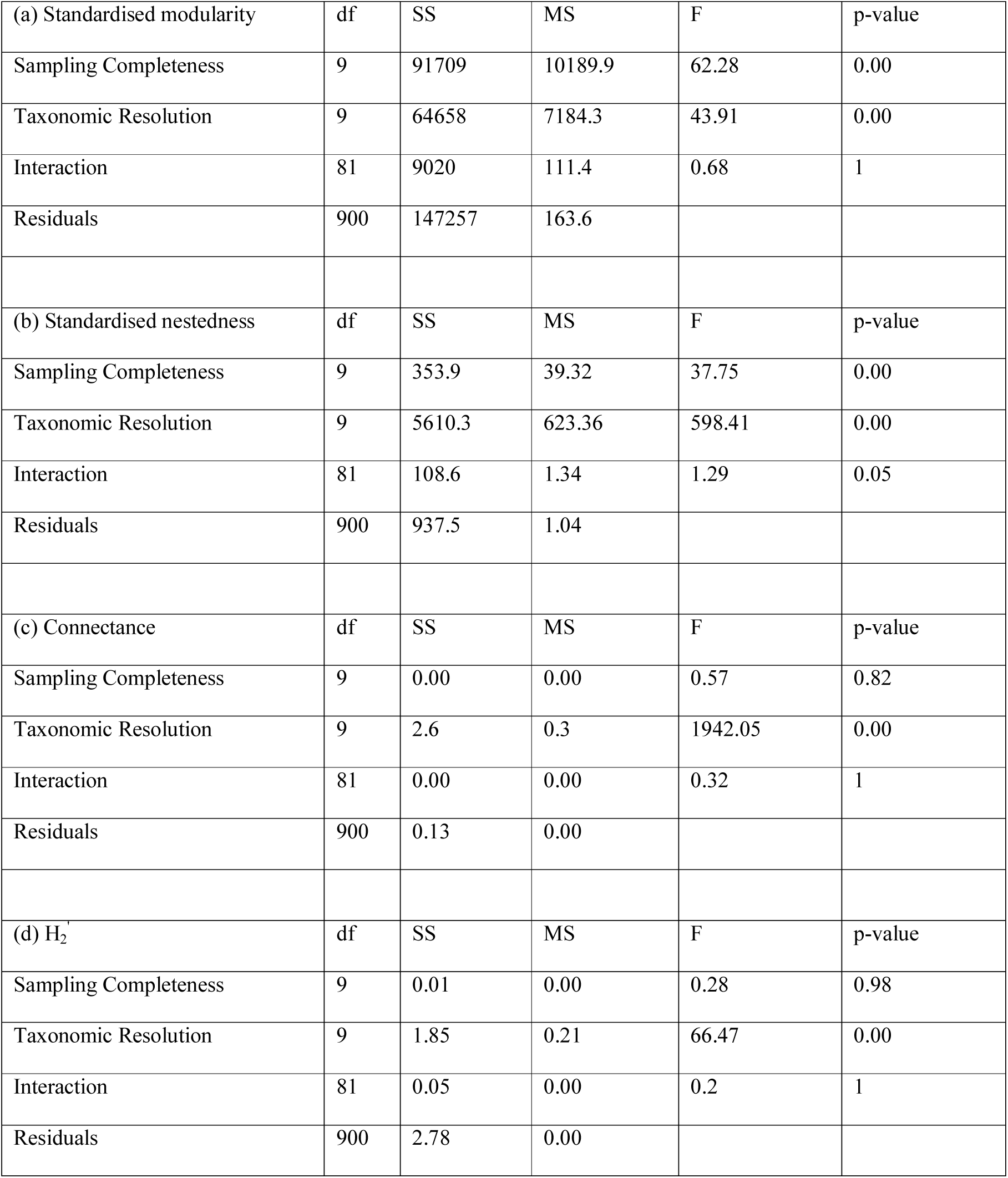
Two-way ANOVA of host sampling completeness and parasite taxonomic resolution for (a) standardised modularity, (b) standardised nestedness, (c) connectance and (d) specialisation (H_2_′).

The four indices were affected by parasite taxonomic resolution (Table 4, Fig. 2, 100% of host sampling completeness). Standardised modularity increased up to 50% of parasite taxonomic resolution in resampled communities, followed by a decline in lower percentages of parasite taxonomic resolution. Estimates of modularity given 70% of parasite taxonomic resolution and below differed from modularity calculated for the full communities (∼0.3 underestimation, Fig. 2a, error bars do not overlap). Both nestedness and connectance increased ∼0.7 and ∼3.2-fold, respectively, as parasite taxonomic resolution decreased. Nestedness estimates at 60% of parasite taxonomic resolution and below differed from nestedness of the full communities (Fig. 2b, error bars do not overlap). We found moderate to strong differences between connectance of the full communities and connectance of the resampled communities at 80% of parasite taxonomic resolution or less (Fig 2c, error bars do not overlap). H_2_′ decreased with parasite taxonomic resolution. We found moderate (∼0.1-fold underestimation) differences between H_2_′ of full communities and resampled communities with 50% or less of parasite taxonomic resolution (Fig. 2d, error bars do not overlap).

The crossed effect of host sampling completeness and parasite taxonomic resolution was additive for both sampling issues for modularity and nestedness, as both effects were significant and no evidence of an interaction was found (Fig. 2, Table 4). Additionally, parasite taxonomic resolution had higher influence on network metrics than host sampling completeness. The standard error of the mean tended to increase with sampling bias, showing proportionally higher variation for connectance than for the other three indices (Fig. 2, Table 4).

Finally, all full and resampled communities were significantly modular. Only in a few severely resampled communities (10% parasite taxonomic resolution capturing less than 50% host sampling completeness) nestedness did not differ from nestedness of 1,000 random assemblages (results not shown). These cases represented Type II errors.

## Discussion

Sampling completeness (representative sample of a community) and taxonomic resolution (accuracy of species assignment) are common sources of uncertainty and bias in community ecology and influence the interpretation of bipartite interactions (Chacoff et al. 2012, Rodrigues and Boscolo 2020). In accordance with our Hypothesis 1, we found that community descriptors are more sensitive to parasite taxonomic resolution than to host sampling completeness. The combination of both sampling issues also resulted in an additive effect for modularity and nestedness (Hypothesis 2, Table 4). Additionally, the descriptors differed in their sensitivity to both sampling issues. This partially concurs with our Hypothesis 3 since modularity was not as robust as in previous studies. Acknowledging that these sampling issues are inevitable to some extent, studies should: (i) avoid applying bipartite network analyses to communities with a low resolution of species identification or low sampling effort; (ii) use the most robust measures to evaluate community structure in communities severely affected by sampling issues (only connectance and H_2_′ were robust to severely under-sampled communities); (iii) pay attention to the conclusions relying on more sensitive metrics (all descriptors were sensitive to taxonomic resolution); and (iv) compare interaction patterns over time and space in communities with comparable and adequate sampling efforts. This is especially true for taxonomic resolution since all descriptors were sensitive to it.

Antagonistic communities are often highly modular (Runghen et al. 2021), most likely due to parasite specialisation on its host resource (Krasnov et al. 2012). We found that modularity was sensitive to both reduction in host sampling completeness and parasite taxonomic resolution. Lower sampling effort left less frequent interactions undetected and eventually decreased resolution in module uniqueness. These results are contrary to those reported in mutualistic communities, in which modularity was found to be a robust descriptor (Rivera-Hutinel et al. 2012, Vizentin-Bugoni et al. 2016). Moreover, an increase in modularity was observed in mutualistic communities with the lowest sampling effort due to increased module identity through removal of between-module interactions (Rivera-Hutinel et al. 2012, Vizentin-Bugoni et al. 2016). A decrease in modularity due to low taxonomic resolution has been found in other plant-insect mutualistic and antagonistic systems (Rodrigues and Boscolo 2020, Renaud et al. 2020). In our study, we found that modularity tended to increase up to a point after which it rapidly decreased in the case of parasite taxonomic resolution, also evident for the crossed effect.

Dallas and Cornelius (2015) found a similar result in their host-parasite co-extinction analyses using in-silico experiments in which they sequentially removed host species first based on species extinction risks and then randomly. These communities with decreasing gradient of host species varied in size and number of interactions, as did our resampled communities which were affected by the crossed effect of both sampling issues. They found that modularity increased up to a critical point and then decreased with both extinction-risk and random removal of host species. However, connectance was robust to random removals but increased with extinction-risk removals in their experiment. Because of the differences between modularity and connectance, they suggested that the behaviour of modularity could be due to either a statistical artifact or that it is a property of complex networks (Dallas & Cornelius, 2015). Here, we suggest that the pattern of initial increase in modularity followed by a decrease may be explained by the way species resolution is lost. Resampled communities from 90% to 60% of parasite taxonomic resolution grouped parasite species with a high overlap in host use. As a result, within-module interactions were strengthened, and modularity increased in these communities. On the other hand, in resampled communities with 20-10% of parasite taxonomic resolution, even parasite species with a low overlap in host use were grouped together. That made host species that actually differ in their parasite communities members of the same module. However, these host species were additionally connected to other modules where the rest of their parasite species were placed. Hence, the loss of parasite taxonomic resolution decreased modularity and made between-module interactions more frequent.

In accordance with our results, nestedness has been shown to be reasonably robust to low sampling completeness in both mutualistic (Nielsen and Bascompte 2007, Vizentin-Bugoni et al. 2016, Fründ et al. 2016) and antagonistic communities (Henriksen et al. 2019). However, as with modularity, previous studies have shown opposite patterns. In earlier studies, nestedness was high in communities with high sampling completeness, and decreased when sampling completeness was lost (Nielsen and Bascompte 2007, Vizentin-Bugoni et al. 2016, Fründ et al. 2016, Henriksen et al. 2019). Interactions in many ecological communities are truly nested (Staniczenko et al. 2013). When such a nested community is under-sampled, one can expect a weaker signature of the nested pattern, or decreasing nestedness with decreasing sampling completeness (Nielsen and Bascompte 2007, Vizentin-Bugoni et al. 2016). However, host-parasite communities are different from other ecological communities and typically show low values of nestedness (Runghen et al. 2021), possibly resulting from coevolution leading to trade-offs in parasite transmission (McQuaid and Britton 2013). The full communities had low values of nestedness (Fig. 2b, negative values or less nested than random assemblages). Hence, low sampling actually opposed the true low nested pattern of antagonistic communities and resulted in increasing nestedness. Moreover, nestedness was not robust since it quickly increased as parasite taxonomic resolution was lost. Along the gradient of taxonomic resolution, parasite species with some shared hosts were grouped. These species appeared as a single generalist parasite species able to infect not only the shared hosts of their foundational parasite species, but also the non-shared hosts. At the same time, specialist parasites remained as separate entities because they were not grouped with any other species. Then, host spectrum of specialist parasites was a subset of the host species used by generalist parasites, or in other words, the network structure became more nested.

Connectance and specialisation (H_2_′) were robust to decreasing host sampling completeness, but not to parasite taxonomic resolution. These descriptors showed opposite patterns, as expected by definition (Table 1). Our results were consistent with previous studies reporting robustness of these metrics to the loss of interactions (Nielsen and Bascompte 2007, Vizentin-Bugoni et al. 2016, Fründ et al. 2016, Henriksen et al. 2019), but not to the loss of species, or the loss of both species and interactions (Dallas and Cornelius 2015, Fründ et al. 2016, Henriksen et al. 2019, Rodrigues and Boscolo 2020, Renaud et al. 2020). The strong dependence of connectance on network size hinders the interpretation of many biological processes (Blüthgen et al. 2006). This pattern was evident along our crossed-effect gradient, where decreasing parasite taxonomic resolution increased the relative contribution of the interactions. H_2_′ was developed as an alternative index of specialisation to overcome the scale dependence issue of connectance (Blüthgen et al. 2006). We found that H_2_′ was sensitive to the loss of parasite taxonomic resolution. However, H_2_′ was only overestimated ∼0.2-fold in the resampled communities with the lowest efforts (10% host sampling completeness × 10% parasite taxonomic resolution), against ∼3-fold in the case of connectance.

We observed that the loss of parasite taxonomic resolution made the descriptor confidence intervals wider. Communities in the gradient of parasite taxonomic resolution vary in size. Large and small communities are known to be biased to different degrees (Shvydka et al. 2018, Henriksen et al. 2019). The full communities represented the largest communities, whereas resampled communities in the lower end of the gradient of parasite taxonomic resolution represented the smallest communities (Table 2b). The later aggregated most of the parasite species with similar infection patterns, losing redundant interactions. Typically, large communities show higher overlap in interaction patterns compared to small communities, where redundancy is low (Henriksen et al. 2019). Therefore, sampling issues are expected to have a limited impact on the network structure of large communities because, if a species is not recorded, the actual infection patterns are nevertheless recorded in redundant interactions. However, inclusion or exclusion of species or interactions produces a greater change among the interaction patterns of small communities. Therefore, the redundancy-size relationship could explain the increasing variance observed in the resampled communities along the parasite taxonomic resolution gradient (Fig. 2, error bars). Nonetheless, severely biased resampled communities were structurally equivalent to full communities (Type II error results) and still useful to categorically describe the structure of ecological communities (e.g., if a community is modular or not), but not the quantitative values of the descriptors (Vanbergen et al. 2017).

The combination of both sampling issues produced an additive effect to modularity and nestedness. Therefore, when both sampling issues coincide in a sampled dataset, this is similarly affected by both sampling issues simultaneously than by the sum of the effects of both sampling issues in isolation. This entails that both sampling issues can be controlled independently. For example, the use of predictive models potentially overcomes the limitation of incomplete interaction richness. These models identify where interactions are most likely to have been missed in a sampled community and eventually include them in the dataset to improve the study of ecological networks (Terry and Lewis 2020). Despite producing greater biases in community descriptors, solutions to taxonomic limitations may not be straightforward. For example, it may require collaborations between ecologists and taxonomists (Poulin and Presswell 2022).

The full communities resembled community and species level descriptors of the natural communities, except for closeness. Earlier studies have evaluated species centrality descriptors, such as closeness, of both parasites (Poulin et al. 2013) and hosts (Dallas et al., 2019). Taxonomic identification at family level was one of the main factors explaining centrality of parasite and host species, suggesting that phylogeny could help in predicting centrality of each species according to their taxonomic affiliation (Poulin et al. 2013). We did not account for the phylogenetic structure of the guilds in the full communities. Instead, each interaction acquired a probability according to the specialisation we represented in the model, but independently of the phylogenetic distance among the members of each guild. The consideration of the phylogenetic structure of parasites and hosts in our model could improve our representation of host-parasite communities and predictions. For example, if parasites infected a range of phylogenetically close hosts, modularity would even be higher because of homogenisation of module composition. Nonetheless, we consider our approach representative of natural communities as most of the community and species level descriptors were effectively captured in the full communities.

Our research shows that studies of communities with low sampling effort and taxonomic resolution may result in wrong conclusions. The implementation of the latest methodological advances in open access software facilitates the use of network analysis in parasitology (Runghen et al. 2021). The increasing availability of host-parasite interaction datasets also favours the comparison or aggregation of communities to address macroecological questions (Doherty et al. 2021). Furthermore, data for some host-parasite communities is already 100 years old. These represent unique case studies to evaluate long-term dynamics and trends at ecosystem level (Carlson et al. 2020) owing to the role of parasites as connectors of all the species they infect throughout their life cycles (Lafferty et al. 2006). However, if communities in such comparative studies notably differ in, or do not include sufficient completeness and species resolution, the conclusions extracted from the network analyses of such data will be of limited use, if not defective.

## Author Contributions

CLB and JJ conceived the ideas; JAB, IBC and VS collected some of the natural communities; AK complied one of the datasets; CLB designed methodology, arranged and analysed data; CLB led the writing of the manuscript. All authors contributed to the drafts and gave final approval for publication.

## Acknowledgements

The authors thank Dr S. Dennis (EAWAG) for his computation assistance. CLB acknowledges the support of an ETH Postdoctoral Fellowship (20-2 FEL-67). VS was funded by the Polish National Agency for Academic Exchange (PPN/ULM/2019/1/00177/U/00001). Data for some natural communities was obtained through funding from the Swiss National Science Foundation (SNSF grant 31003A_169211) to IBC.

## Supporting information

Llopis-Belenguer, C., J. A. Balbuena, I. Blasco-Costa, A. Karvonen, V. Sarabeev, J. Jokela. Sensitivity of bipartite network analyses to incomplete sampling and taxonomic uncertainty.

## Appendix S1: Data sources

- Full communities: Replication code and data for these analyses is available in: Llopis-Belenguer, C., J. A. Balbuena, I. Blasco-Costa, A. Karvonen, V. Sarabeev, J. Jokela. Replication code and data for: Sensitivity of bipartite network analyses to incomplete sampling and taxonomic uncertainty. *Zenodo*, DOI### (to be published in Zenodo after acceptance)
- Natural communities (retrieved from http://www.ecologia.ib.usp.br/iwdb/): Arai, H. P., and D. R. Mudry. 1983. Protozoan and Metazoan Parasites of Fishes from the Headwaters of the Parsnip and McGregor Rivers, British Columbia: A Study of Possible Parasite Transfaunations. Canadian Journal of Fisheries and Aquatic Sciences 40:1676–1684. Arthur, J. R., L. Margolis, and H. P. Arai. 1976. Parasites of Fishes of Aishihik and Stevens Lakes, Yukon Territory, and Potential Consequences of Their Interlake Transfer Through a Proposed Water Diversion for Hydroelectrical Purposes. Journal of the Fisheries Research Board of Canada 33:2489–2499. Chinniah, V. C., and W. Threlfall. 1978. Metazoan parasites of fish from the Smallwood Reservoir, Labrador, Canada. Journal of Fish Biology 13:203–213. Leong, T. S., and J. C. Holmest. 1981. Communities of metazoan parasites in open water fishes of Cold Lake, Alberta. Journal of Fish Biology 18:693–713.
- Natural communities and Host sampling completeness (data in Valtonen et al. (2001)): Llopis-Belenguer, C. et al. Parasite communities of fishes from Northeastern Baltic Sea (data in Valtonen et al., 2001). *Zenodo*, DOI### (to be published in Zenodo after acceptance)
- Host sampling completeness Llopis-Belenguer, C., 2019. Replication Data for: Native and invasive hosts play different roles in host-parasite networks. *Harvard Dataverse*, https://doi.org/10.7910/DVN/IWIKOL Llopis-Belenguer, C., J.A. Balbuena, K. Lange, F. de Bello, I. Blasco-Costa. 2018. Data from: Unveiling hidden diversity: towards a functional trait framework for parasites. *Harvard Dataverse*, https://doi.org/10.7910/DVN/RX3R2X Llopis-Belenguer, C. et al. Parasite communities of *Coregonus* spp. from Swiss and Norwegian Lakes. *Zenodo*, DOI### (to be published in Zenodo after acceptance)

## Appendix S2: Fig. S1

**Fig. S1.**
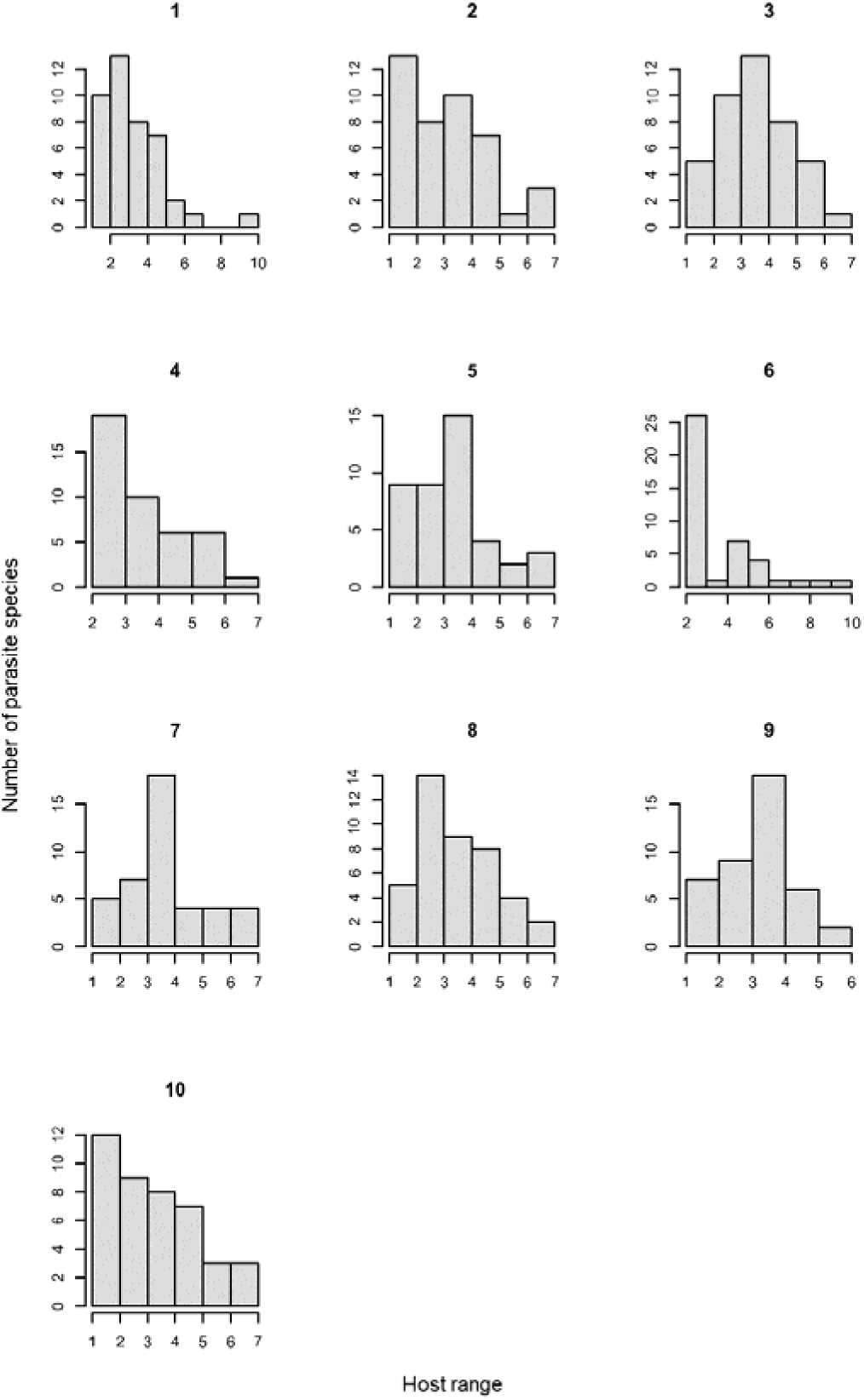
Host range distribution of the parasite species in the 10-replicate full communities. Bar plots show the number of parasite species that infect a given number of host species. Almost all parasite species infect less than half of the host species (nhost=13) due to the high specialisation parameter (specpar=55). The mean intensity of each parasite species in a given host species was not considered in these plots, but presence-absence data.

## References

1. Almeida-Neto, M., and W. Ulrich. 2011. A straightforward computational approach for measuring nestedness using quantitative matrices. Environmental Modelling & Software 26:173–178.

2. Beckett, S. J. 2016. Improved community detection in weighted bipartite networks. Royal Society open science 3:140536.

3. Bellekom, B., T. D. Hackett, and O. T. Lewis. 2021. A Network Perspective on the Vectoring of Human Disease. Trends in Parasitology 37:391–400.

4. Blüthgen, N. 2010. Why network analysis is often disconnected from community ecology: A critique and an ecologist’s guide. Basic and Applied Ecology 11:185–195.

5. Blüthgen, N., F. Menzel, and N. Blüthgen. 2006. Measuring specialization in species interaction networks. BMC Ecology 6:9.

6. Cardoso, T. D. S., C. S. Andreazzi, A. Maldonado Junior, and R. Gentile. 2021. Functional traits shape small mammal-helminth network: patterns and processes in species interactions. Parasitology 148:947–955.

7. Carlson, C. J., S. Hopkins, K. C. Bell, J. Doña, S. S. Godfrey, M. L. Kwak, K. D. Lafferty, M. L. Moir, K. A. Speer, G. Strona, M. Torchin, and C. L. Wood. 2020. A global parasite conservation plan. Biological Conservation 250:108596.

8. Chacoff, N. P., D. P. Vázquez, S. B. Lomáscolo, E. L. Stevani, J. Dorado, and B. Padrón. 2012. Evaluating sampling completeness in a desert plant–pollinator network. Journal of Animal Ecology 81:190–200.

9. Dallas, T. A., B. A. Han, C. L. Nunn, A. W. Park, P. R. Stephens, and J. M. Drake. 2019. Host traits associated with species roles in parasite sharing networks. Oikos 128:23–32.

10. Dallas, T., and E. Cornelius. 2015. Co-extinction in a host-parasite network: identifying key hosts for network stability. Scientific Reports 5:13185.

11. Delmas, E., M. Besson, M. H. Brice, L. A. Burkle, G. V. Dalla Riva, M. J. Fortin, D. Gravel, P. R. Guimarães, D. H. Hembry, E. A. Newman, J. M. Olesen, M. M. Pires, J. D. Yeakel, and T. Poisot. 2019. Analysing ecological networks of species interactions. Biological Reviews 94:16–36.

12. Doherty, J. F., X. Chai, L. E. Cope, D. de Angeli Dutra, M. Milotic, S. Ni, E. Park, and A. Filion. 2021. The rise of big data in disease ecology. Trends in Parasitology 37:1034– 1037.

13. Dormann, C. F. 2011. How to be a specialist? Quantifying specialisation in pollination networks. Network Biology 1:1–20.

14. Dormann, C. F., J. Fründ, and H. M. Schaefer. 2017. Identifying Causes of Patterns in Ecological Networks: Opportunities and Limitations. Annual Review of Ecology, Evolution, and Systematics 48:559–584.

15. Ferguson, K. I., and P. Stiling. 1996. Non-additive effects of multiple natural enemies on aphid populations. Oecologia 108:375–379.

16. Frankham, R., J. D. Ballou, K. Ralls, M. D. B. Eldridge, M. R. Dudash, C. B. Fenster, R. C. Lacy, and P. Sunnucks. 2017. Population fragmentation causes inadequate gene flow and increases extinction risk. Page Genetic Management of Fragmented Animal and Plant Populations. Oxford University Press.

17. Fründ, J., K. S. McCann, and N. M. Williams. 2016. Sampling bias is a challenge for quantifying specialization and network structure: lessons from a quantitative niche model. Oikos 125:502–513.

18. Gotelli, N. J., and K. Rohde. 2002. Co-occurrence of ectoparasites of marine fishes: A null model analysis. Ecology Letters 5:86–94.

19. Guimarães Jr., P. R. 2020. The Structure of Ecological Networks Across Levels of Organization. Annual Review of Ecology, Evolution, and Systematics 51:433–460.

20. Henriksen, M. V, D. G. Chapple, S. L. Chown, and M. A. McGeoch. 2019. The effect of network size and sampling completeness in depauperate networks. Journal of Animal Ecology 88:211–222.

21. Isaac, N. J. B., J. Mallet, and G. M. Mace. 2004. Taxonomic inflation: its influence on macroecology and conservation. Trends in Ecology & Evolution 19:464–469.

22. Kozłowski, J., and A. T. Gawelczyk. 2002. Why are species’ body size distributions usually skewed to the right? Functional Ecology 16:419–432.

23. Krasnov, B. R., M. A. Fortuna, D. Mouillot, I. S. Khokhlova, G. I. Senbrot, and R. Poulin. 2012. Phylogenetic Signal in Module Composition and Species Connectivity in Compartmentalized Host-Parasite Networks. The American Naturalist 179:501–511.

24. Lafferty, K. D., A. P. Dobson, and A. M. Kuris. 2006. Parasites dominate food web links. Proceedings of the National Academy of Sciences 103:11211–11216.

25. Llopis-Belenguer, C., S. Pavoine, I. Blasco-Costa, and J. A. Balbuena. 2020. Assembly rules of helminth parasite communities in grey mullets: combining components of diversity. International Journal for Parasitology 50:1089–1098.

26. Maechler, M., P. Rousseeuw, A. Struyf, M. Hubert, and R. K. Horniks. 2021. cluster: Cluster Analysis Basics and Extensions.

27. Martín González, A. M., B. Dalsgaard, and J. M. Olesen. 2010. Centrality measures and the importance of generalist species in pollination networks. Ecologssssssssssical Complexity 7:36– 43.

28. Martinez, N. D. 1991. Artifacts or attributes? Effects of resolution on the Little Rock Lake food web. Ecological Monographs 61:367–392.

29. McQuaid, C. F., and N. F. Britton. 2013. Host-parasite nestedness: A result of co-evolving trait-values. Ecological Complexity 13:53–59.

30. Morand, S., and B. Krasnov. 2008. Why apply ecological laws to epidemiology? Trends in Parasitology 24:304–309.

31. Morris, R. J., S. Gripenberg, O. T. Lewis, and T. Roslin. 2014. Antagonistic interaction networks are structured independently of latitude and host guild. Ecology Letters 17:340–349.

32. Newman, M. E. J. 2001. Scientific collaboration networks. II. Shortest paths, weighted networks, and centrality. Physical Review E 64:016132.

33. Nielsen, A., and J. Bascompte. 2007. Ecological networks, nestedness and sampling effort. Journal of Ecology 95:1134–1141.

34. Pavoine, S., A.-B. Dufour, and D. Chessel. 2004. From dissimilarities among species to dissimilarities among communities: a double principal coordinate analysis. Journal of Theoretical Biology 228:523–537.

35. Poisot, T., D. B. Stouffer, and D. Gravel. 2015. Beyond species: why ecological interaction networks vary through space and time. Oikos 124:243–251.

36. Poulin, R. 1998. Comparison of Three Estimators of Species Richness in Parasite Component Communities. The Journal of Parasitology 84:485–490.

37. Poulin, R. 2007. Evolutionary Ecology of Parasites. Princeton University Press.

38. Poulin, R. 2010. Network analysis shining light on parasite ecology and diversity. Trends in Parasitology 26:492–498.

39. Poulin, R., B. R. Krasnov, S. Pilosof, and D. W. Thieltges. 2013. Phylogeny determines the role of helminth parasites in intertidal food webs. Journal of Animal Ecology 82:1265– 1275.

40. Poulin, R., and T. L. F. Leung. 2010. Taxonomic resolution in parasite community studies: are things getting worse? Parasitology 137:1967–1973.

41. Poulin, R., and S. Morand. 1997. Parasite body size distributions: interpreting patterns of skewness. International Journal for Parasitology 27:959–964.

42. Poulin, R., and B. Presswell. 2022. Is parasite taxonomy really in trouble? A quantitative analysis. International Journal for Parasitology 52:469–474.

43. R Core Team. 2021. R: A language and environment for statistical computing. R Foundation for Statistical Computing.

44. Renaud, E., E. Baudry, and C. Bessa-Gomes. 2020. Influence of taxonomic resolution on mutualistic network properties. Ecology and Evolution 10:3248–3259.

45. Ricotta, C., and L. Szeidl. 2009. Diversity partitioning of Rao’s quadratic entropy. Theoretical Population Biology 76:299–302.

46. Rivera-Hutinel, A., R. O. Bustamante, V. H. Marín, and R. Medel. 2012. Effects of sampling completeness on the structure of plant–pollinator networks. Ecology 93:1593–1603.

47. Rodrigues, B. N., and D. Boscolo. 2020. Do bipartite binary antagonistic and mutualistic networks have different responses to the taxonomic resolution of nodes? Ecological Entomology 45:709–717.

48. Runghen, R., R. Poulin, C. Monlleó-Borrull, and C. Llopis-Belenguer. 2021. Network Analysis: Ten Years Shining Light on Host–Parasite Interactions. Trends in Parasitology 37:445–455.

49. Shvydka, S., V. Sarabeev, V. D. Estruch, and C. Cadarso-Suárez. 2018. Optimum sample size to estimate mean parasite abundance in fish parasite surveys. Helminthologia (Poland) 55:52–59.

50. Staniczenko, P. P. A., J. C. Kopp, and S. Allesina. 2013. The ghost of nestedness in ecological networks. Nature Communications 4:1391.

51. Thioulouse, J., S. Dray, A.-B. Dufour, A. Siberchicot, T. Jombart, and S. Pavoine. 2018. Multivariate Analysis of Ecological Data with ade4. Page Multivariate Analysis of Ecological Data with ade4. Springer New York, New York, NY.

52. Thompson, R. M., and C. R. Townsend. 2000. Is resolution the solution?: the effect of taxonomic resolution on the calculated properties of three stream food webs. Freshwater Biology 44:413–422.

53. Tylianakis, J. M., T. Tscharntke, and O. T. Lewis. 2007. Habitat modification alters the structure of tropical host–parasitoid food webs. Nature 445:202–205.

54. Valtonen, E. T., K. Pulkkinen, R. Poulin, and M. Julkunen. 2001. The structure of parasite component communities in brackish water fishes of the northeastern Baltic Sea. Parasitology 122:471–481.

55. Vanbergen, A. J., B. A. Woodcock, M. S. Heard, and D. S. Chapman. 2017. Network size, structure and mutualism dependence affect the propensity for plant–pollinator extinction cascades. Functional Ecology 31:1285–1293.

56. Vázquez, D. P., C. J. Melián, N. M. Williams, N. Blüthgen, B. R. Krasnov, and R. Poulin. 2007. Species abundance and asymmetric interaction strength in ecological networks. Oikos 116:1120–1127.

57. Vizentin-Bugoni, J., P. K. Maruyama, V. J. Debastiani, L. da S. Duarte, B. Dalsgaard, and M. Sazima. 2016. Influences of sampling effort on detected patterns and structuring processes of a Neotropical plant–hummingbird network. Journal of Animal Ecology 85:262–272.

